# Excessive ERK-dependent synaptic clustering drives enhanced motor learning in the MECP2 duplication syndrome mouse model of autism

**DOI:** 10.1101/100875

**Authors:** Ryan Thomas Ash, Shelly Alexandra Buffington, Jiyoung Park, Mauro Costa-Mattioli, Huda Yaya Zoghbi, Stelios Manolis Smirnakis

## Abstract

Autism-associated genetic mutations may produce altered learning abilities by perturbing the balance between stability and plasticity of synaptic connections in the brain. Here we report an increase in the stabilization of dendritic spines formed during repetitive motor learning in the mouse model of *MECP2*-duplication syndrome, a high-penetrance form of syndromic autism. This increased stabilization is mediated entirely by spines that form cooperatively in clusters. The number of clusters formed and stabilized predicts the mutant’s enhanced motor learning and memory phenotype, reminiscent of savant-like behaviors occasionally associated with autism.

The ERK signaling pathway, which promotes cooperative plasticity between spines, was found to be hyperactive in *MECP2*-duplication motor cortex specifically after training. Inhibition of ERK signaling normalizes clustered spine stabilization and rescues motor learning behavior in mutants. We conclude that learning-associated dendritic spine clustering stabilized by hyperactive ERK signaling drives abnormal motor learning and memory consolidation in this model of syndromic autism.

## Introduction

It has been proposed that phenotypes of autism spectrum disorder arise from an abnormal imbalance between the stability and plasticity of synaptic connections in the brain (Ramocki et al., 2008). Such an imbalance may potentially contribute to the rigid, restricted behavioral repertoire, insistence on sameness, and at times savant-like behavior seen in autism^1,2^. Rebalancing synaptic stability and plasticity could provide a therapeutic avenue to promote behavioral flexibility in patients. The goal of the present study was to test this proposal by linking changes in synaptic plasticity to a robust learning phenotype in an animal model of autism *in vivo*.

Methyl-CpG-binding protein 2 (MeCP2) is a transcriptional regulator that contributes to the maintenance of neural circuit homeostasis through the activity-dependent modulation of gene expression and splicing^3^. Loss of function mutations in *MECP2* cause Rett Syndrome, a neurodevelopmental disorder affecting females^4,5^. Mice engineered to overexpress *MECP2* at twice normal levels exhibit a neurological phenotype that falls squarely within the spectrum of autism^6^. They exhibit social avoidance, stereotypies, behavioral inflexibility as well as initially enhanced motor learning and memory that progresses to motor dysfunction, spasticity and epilepsy^6–8^. Patients with genomic duplication of *MECP2,* identified soon after the description of *MECP2* duplication syndrome in mice, demonstrate many similar features highly characteristic of autism^9–11^. The MECP2-duplication mouse is therefore a valuable tool for studying neural circuit phenotypes in autism^12^.

Enhanced motor learning and memory with repetitive training on the rotarod task is a phenotype shared across several mouse models of autism, including MECP2-duplication mice, and studied as a model for consolidation of repetitive motor behaviors^6,13–16^. Interestingly these mouse models also share an upregulation in the spontaneous turnover of pyramidal neuron dendritic spines^17–20^, suggestive of an abnormal balance between synaptic stability and plasticity. A change in *learning-associated* dendritic spine plasticity in these mice, however, has not been previously assessed.

Repetitive motor training is known to induce the formation of dendritic spines (synapses) onto the apical dendrites of corticospinal neurons in normal mouse primary motor cortex^21,22^. The number of spines stabilized correlates with performance, and experimental ablation of stabilized learning-associated spines disrupts the motor memory associated with those spines^23^, suggesting that learning-associated spine stabilization represents a durable structural correlate of procedural skill memory. It is an important open question whether the process of learning-associated dendritic spine stabilization is abnormal in autism.

In what follows, we provide evidence that the abnormal repetitive motor learning phenotype in *MECP2*-duplication mice arises from excessive cooperative stabilization between neighboring synaptic connections formed during learning. Excessive stabilization of synaptic clusters, but not of isolated spines, predicts enhanced motor learning in mutant animals. Both excessive synaptic clustering and enhanced motor learning are associated with a training-specific increase in ERK signaling in motor cortex, and can be reversed by pharmacological normalization of ERK signaling. These data identify a structural neural circuit abnormality linking a genetic mutation and cell-signaling pathway to a quantifiable learning phenotype in a mouse model of autism, in vivo.

## Results

### Increased learning-associated dendritic spine stabilization and enhanced repetitive motor learning in *MECP2*-duplication mice

We employed in vivo 2-photon microscopy^24^ in primary motor cortex (M1) to relate the stability of dendritic spines formed during motor learning^21^ to the MECP2-duplication mouse’s enhanced learning on the rotarod task (Fig. 1a,b), a repetitive motor learning paradigm^13
14
25^. Apical dendrites from GFP-expressing^26^ complex-tufted L5 pyramidal neurons in area M1, primarily corticospinal neurons^27^, were targeted for imaging (Fig. 1a, Supplementary Movie 1)^24^. Spine analysis was performed on terminal dendritic branches of the apical tuft of these neurons. Imaging, training, and analysis were performed blind to genotype (see Methods), and correct targeting to area M1 was confirmed by electrical microstimulation after the final imaging session (Supplementary Fig. 1b)^28^. We first identified baseline (pre-training) spines, then trained the animals on the rotarod for four days (Fig. 1b). On the fifth day, we imaged the dendrites again to identify new spines formed with training (learning-associated spines). We found that approximately 30% more spines were formed with training in MECP2-duplication mice (Fig. 1c, orange) vs. wild-type littermates (WT, black) (Fig. 1c, 4.1±0.2 vs. 2.9±0.3 spines/100μm, *P*=0.02, n=10 animals per genotype, Mann-Whitney U test). Following four days of rest (the time frame of spine stabilization^29^), dendrites were once again imaged to identify the learning-associated spines that stabilized. Almost twice as many learning-associated spines were stabilized in *MECP2*-duplication mice compared to controls (Fig. 1d, 1.5±0.1 vs. 0.8±0.2 spines/100μm, *P*=0.02; n=10 animals per genotype; Mann-Whitney U test).

**Figure 1.**
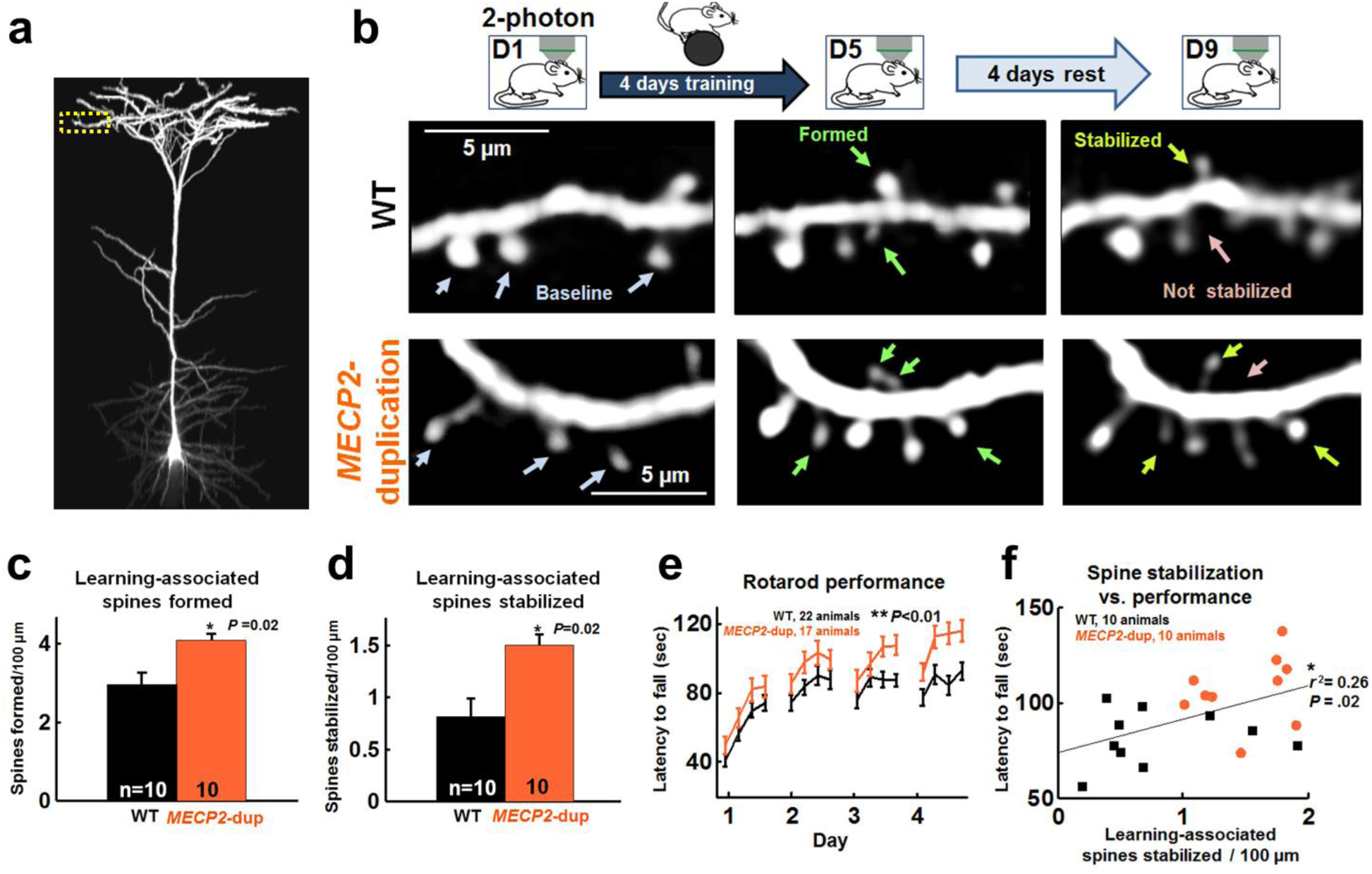
Learning-associated spine stabilization correlates with enhanced motor learning in *MECP2*-duplication mice. **a**, The dendritic tree of a GFP-labeled L5 complex-tufted pyramidal neuron, i.e. corticospinal neuron, in area M1 imaged by in vivo 2-photon microscopy. Apical tuft terminal dendrites are targeted for time-lapse imaging (yellow box). **b**, Experiment paradigm: Sample images of dendritic segments imaged before training (left), following 4 days of training (middle), and 4 days after the end of training (right). *Top:* Wild-type controls. *Bottom: MECP2*-duplication animals. White arrows point to spines present at baseline, green to spines formed during learning, light-green to stabilized spines, and pink to non-stabilized spines. **c**, Motor-learning-associated spines formed per 100μm in terminal dendritic branches of *MECP2*-duplication animals (orange, n=10 animals, 300 spines formed, 82 dendritic branches) and WT littermate controls (black, n=10 animals, 276 spines formed, 99 dendritic branches). * *P* = 0.014, Z(18)=−2.45, Mann-Whitney U test across animals. **d**, Learning-associated spines stabilized per 100μm in each genotype. * *P* = 0.017. **e**, Mean per-trial rotarod performance (time spent on the accelerating rotarod before falling) across animals in *MECP2*-duplication animals (orange, n=17) and WT controls (black, n=22)^6^. **, *P* = 0.009, effect of genotype, F(1,37)=7.6, repeated measures ANOVA. Effect of time: *P* < 0.0001, F(15,37)=29.8; Interaction: *P* =0.47, F(15,37)=0.99. **f**, Scatter plot of learning associated spines that stabilized versus rotarod performance per animal, in *MECP2*-duplication mice (orange circles) and WT controls (black squares). * *P* < 0.05, r^2^=0.26, n=20 mice pooled across genotypes, Pearson correlation, student’s t test. Error bars indicate mean ± s.e.m.

*MECP2*-duplication mice performed significantly better on the rotarod as has been previously reported^6^ (*P*=0.01; WT: n=22 animals, *MECP2*-duplication: n=17; repeated-measures ANOVA) (Fig. 1e, Supplementary Movie 2). The median best per-day performance was 117±5 sec for *MECP2*-duplication mice and 94±4 sec for controls (Supplementary Fig. 1c, P=10^−4^, t test across animals). Learning-associated spine stabilization correlated with rotarod performance in individual animals^21^ (r^2^= 0.26, P=0.02, Pearson correlation, student’s t test), suggesting that a bias toward learning-associated synapse stability contributes to the mutant’s motor learning phenotype (Fig. 1f). In contrast to previous results^21,30^, WT control rotarod performance did not by itself correlate with spine stabilization (r^2^= 0.07, *P*=0.46, n=10, Pearson correlation, student’s t test), possibly due to differences in rotarod training paradigm (4 trials per day for 4 days in this study vs. 15–20 trials per day for 2 days in previous studies). Prior to training, one-day spontaneous spine formation and elimination in motor cortex L5 apical dendrites was comparable between *MECP2*-duplication mice and WT littermates (Supplementary Fig. 1d), indicating that the differences we measure here between mutants and WT manifest specifically with training. Elimination of baseline spines was not significantly different in mutants (Supplementary Fig. 1e,f), and baseline spine elimination did not predict rotarod performance (Supplementary Fig. 1g, r^2^= 0.04, *P*=0.4). New spine formation in the 4 days following the end of training, terminal dendritic branch lengths, and spine densities in apical tufts of corticospinal M1 neurons were also similar between 4-month-old *MECP2*-duplication mice and WT controls (Supplementary Fig. 1h-j).

### Increased stabilization of dendritic spine clusters following training in *MECP2*- duplication mice

We next examined the spatial distribution of the spines forming the new procedural memory trace^31,32^. As previously reported in normal mice undergoing repeated motor training^33^, we observed that in *MECP2*-duplication mice learning-associated spines were often stabilized in pairs or triplets along the dendrite (Fig. 1b, bottom right panel, Supplementary Fig. 2). We binned learning-associated stabilized spines by their proximity to other learning-associated spines and detected a dramatic increase in synaptic clustering in *MECP2*-duplication mice that was not present in WT littermates (Fig. 2a). About 3 times as many spines were stabilized within 5 μm of another learning-associated spine in *MECP2*-duplication mice compared to WT controls (6.4±0.7 vs. 2.3±0.7 spines/1000μm, *P*<0.0001, n=10 animals per genotype, ANOVA with Tukey correction for multiple comparisons), and almost twice as many spines were stabilized within 5–10 μm of another new spine (3.0±0.5 vs. 1.6±0.5 spines/1000μm, though this difference did not reach significance). Beyond 10 μm interspine distance, similar numbers of spines were stabilized in both genotypes, indicating that the increased spine stabilization observed in *MECP2*-duplication mice (Fig. 1d) is mediated almost exclusively through excessive stabilization of dendritic spine clusters (Fig. 2a; Supplementary Fig. 3a). Simulations showed that increased clustered spine stability was not a byproduct of the overall increase in learning-associated spine formation or stabilization observed in mutants (Supplementary Fig. 3a, dotted lines). Non-stabilized spines occurred in clusters at similar rates between WT mice and *MECP2*-duplication mice (Fig. 2b, Supplementary Fig. 3b,d). The distance from each stabilized spine to the nearest non-stabilized learning-associated spine (Supplementary Fig. 3b) or baseline spine (Supplementary Fig. 3c) did not differ significantly between *MECP2*-duplication mice and controls (Fisher exact test), and baseline spines also demonstrated spatial distributions similar to WT (Supplementary Fig. 3e, Fisher exact test).

**Figure 2.**
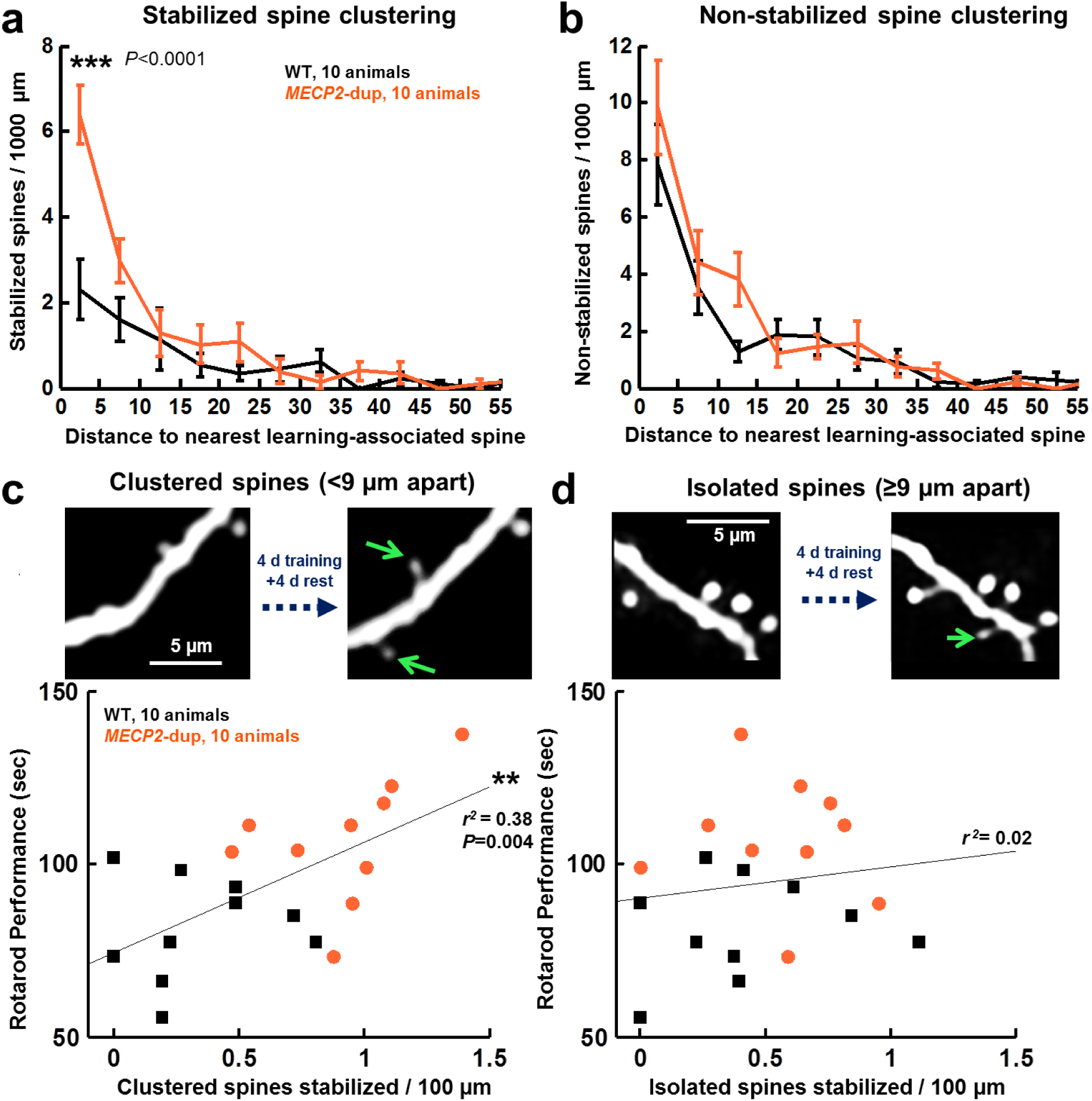
Increased stabilization of dendritic spine clusters in *MECP2*-duplication mice. **a**, Mean number of learning-associated spines that stabilized per 1000μm, binned by distance to the nearest neighboring learning-associated spine. Bin size = 5μm. *** *P* = 10^−7^, 2-way ANOVA with Tukey correction for multiple comparisons, Fgenotype(1,19)=14.3, F_clusterbin_(12,19)=22.6, F_genotype × clusterbin_(12,19)=5.13, n=10 animals per genotype. **b**, Mean number of learning-associated spines that formed but didn’t stabilize per 1000μm, binned by distance to the nearest neighboring learning-associated spine, F_genotype_(1,19)=2.2, F_clusterbin_(12,19)=29.3, F_genotype × clusterbin_(12,19)=1.1, n=10 animals per genotype. **c**, Stabilization of learning-associated dendritic spine clusters (<9μm to the nearest neighboring learning-associated spine) correlates with enhanced rotarod performance in *MECP2*-duplication mice (orange dots) and WT controls (black squares). ** *P* = 0.004, r^2^=0.38, n=20 mice pooled across genotypes, Pearson correlation, student’s t test. Supplementary Fig. 4A and Methods explain how the 9μm threshold was chosen. **d**, *Top:* example of one isolated learning-associated stabilized spine. Stabilization of isolated learning-associated spines (≥9μm to the nearest neighboring learning-associated spine) does not correlate with rotarod performance (r^2^=0.02, P=0.55), suggesting that learning associated spine clustering is a better predictor of behavioral performance. Error bars indicate mean ± s.e.m.

To study the relationship between clustered-spine stabilization and motor performance in individual mice, we categorized each stabilized spine as clustered (<9 microns to nearest neighboring learning-associated spine, Fig. 2c) or isolated (≥9 microns to nearest neighboring learning-associated spine, Fig. 2d). Nine microns was chosen as a distance threshold for defining clusters because the difference in clustered-spine stabilization between mutants and WT plateaued at this threshold (Supplementary Fig. 4a, see Methods). Clustered learning-associated spines were almost twice as likely to be stabilized in *MECP2*-duplication mice (41±3% of clustered spines) compared to WT (22±4% of clustered spines, *P*= 0.01, Mann-Whitney U test, data not shown). Remarkably, the number of stabilized dendritic spine clusters correlated well with enhanced rotarod performance, while the number of stabilized isolated spines did not (Fig. 2c,d, clustered spines: *r*^2^ = 0.38, *P*=0.004; isolated spines: *r*^2^ = 0.02, *P*=0.55; student’s t distribution). Clustered spine stabilization was increased in *MECP2*- duplication mice even when comparisons were limited to mice that were matched for time spent on the rotarod (Supplementary Fig. 4b); increased clustered spine stabilization is therefore not a simple consequence of increased time spent on the rotarod. These results suggest that increased *clustered-spine* stabilization occurs with training in *MECP2*-duplication mice and promotes their enhanced learning.

### Enhanced motor learning in *MECP2*-duplication mice is ERK dependent

The increased clustered-spine stability in the ~9–10 μm range we observe here is strikingly similar to the range of a known ERK-dependent form of clustered-spine plasticity^34–36^. A simplified schematic of ERK signaling in clustered-spine plasticity is illustrated in Fig. 3a^34,37^. Given that several genes in the ERK pathway are associated with autism^38^ (asterisks in Fig. 3a) and that *MECP2*-duplication mice overexpress several genes in this pathway^39^ (notably *Bdnf, Nmdar1,* and *Ras,* underlined in Fig. 3a), we hypothesized that hyperactive ERK signaling may contribute to the learning and plasticity phenotypes observed in our animals. To directly test if ERK signaling is increased during motor learning in *MECP2*-duplication mice, we performed Western blot analyses on M1 protein extracts isolated from mutant mice and WT littermates at baseline and following rotarod training (Fig. 3b, Supplementary Fig. 5a). We found that a marked hyper-phosphorylation of the MAP kinase ERK1/2 (T202/Y204) in *MECP2*- duplication mouse area M1 occurred specifically with rotarod training (Fig. 3b, P=0.004, n=10 WT, 12 *MECP2*-duplication, 2-tailed unpaired t-test). Remarkably, the phosphorylation state of ERK was not significantly altered in mutant M1 lysates prior to training (P=0.87, n=6 mice per genotype, 2-tailed unpaired t-test)^40^. Note that in M1, ERK phosphorylation did not correlate with 1^st^-day latency to fall from the rotarod (Supplementary Fig. 5b, r=−0.17, p= 0.53), so increased ERK activation in mutants could not be explained solely by increased time spent on the rotarod.

**Figure 3.**
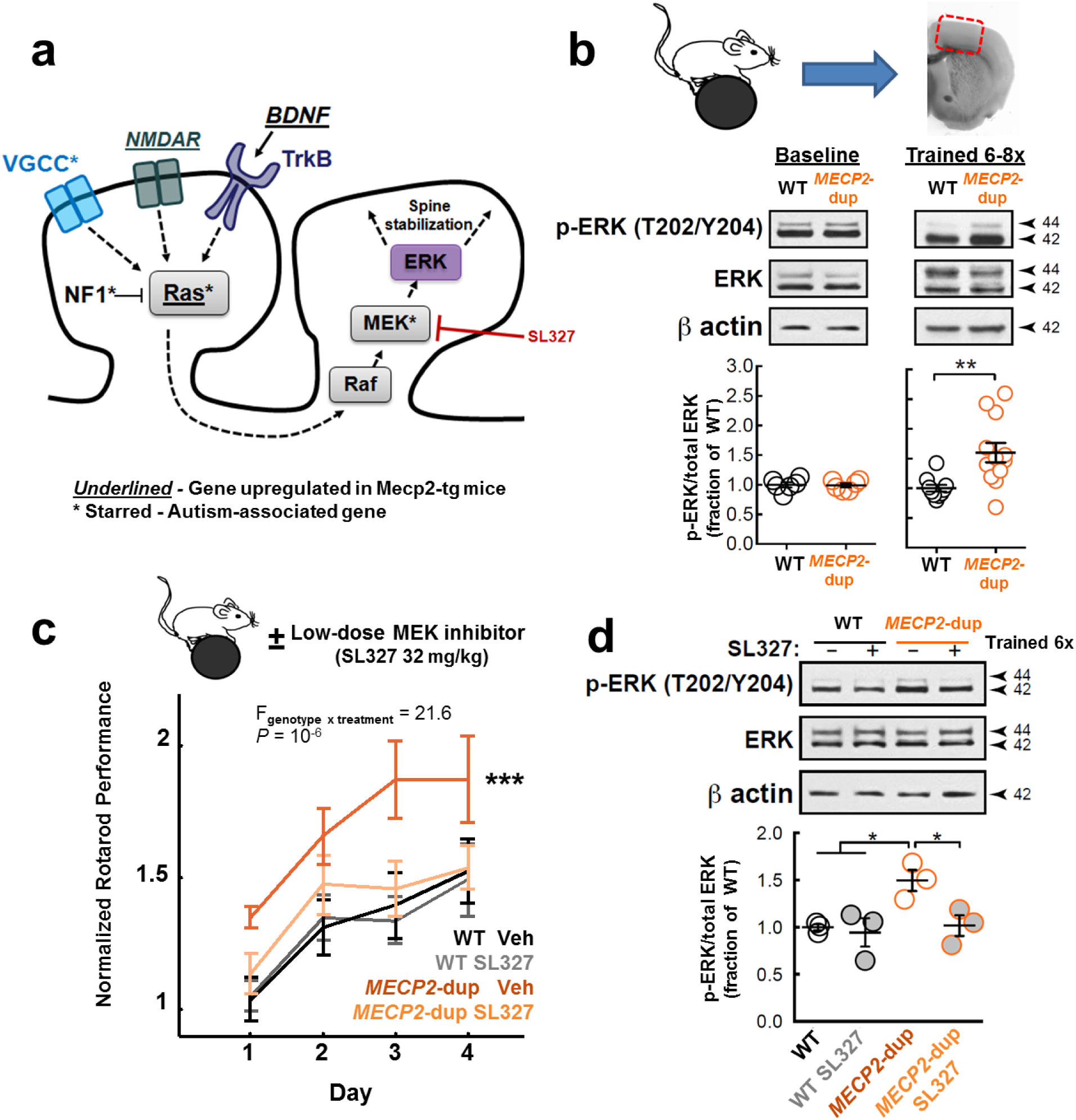
Enhanced motor learning in *MECP2*-duplication mice is ERK dependent. **a**, A simplified schema of the Ras-MAPK signaling pathway and how it is hypothesized to contribute to clustered-spine stabilization^34^. Genes transcriptionally upregulated in MECP2-overexpressing mice^39^ are underlined. Known autism-associated genes are denoted with an asterisk^38^. **b**, Rotarod training (6–8 trials) induces enhanced Ras-MAPK signaling (ERK phosphorylation) in *MECP2*-duplication mice relative to WT littermates, despite equivalent baseline levels prior to training. Representative Western blots and densitometric quantification of ERK activation (p-ERK T202/Y204 / total ERK immunoreactivity; 42 and 44 kD bands summed in quantification) before (baseline; n=6 mice/genotype; unpaired 2-sided t-test: *P*=0.9, t(10)=0.17) and after training (n=10 WT and 12 *MECP2*-duplication mice; unpaired 2-sided t-test: ** *P*=0.004, t(20)=3.2). Data are presented as fraction of the WT mean. **c**, The MEK inhibitor SL327 normalizes rotarod performance in *MECP2*-duplication mice. Mean±sem of the peak performance on each day plotted for vehicle-treated *MECP2*-duplication (dark orange, n=8 mice), vehicle-treated WT (black, n=7), SL327-treated *MECP2*-duplication (pale orange, n=9), and SL327-treated WT (gray, n=9) mice. Data are presented per day and normalized by the mean first-day performance of WT littermates for illustration purposes. *** *P* = 10^−6^, genotype x drug interaction, F_genotype x treatment_(1,1,29) = 21.6; F_genotype_(1,29)=6.4, *P* = 0.01; F_treatment_(1,29)=4.9, p=0.027; F_trial_(15,29)=15.8, *P* = 10^−6^; mixed effects repeated-measures ANOVA. Statistical analysis was performed on raw per-trial rotarod performance values. Litter was included as a variable to control for across-litter variability in performance. **d**, Consistent with a role for elevated ERK signaling in the mutant’s enhanced motor learning phenotype, 32mg/kg SL327 treatment blocked the training-dependent increase in M1 ERK phosphorylation in *MECP2*-duplication mice. Representative Western blots and densitometric quantification of ERK activation in vehicle-versus SL327-treated WT and *MECP2*-duplication mice (n=3 mice/genotype/treatment group. * P < 0.05, F_genotype_(1,8)= 6.7, F_treatment_(1,8)=5.9, F_interaction_(1,1,8)=3.7, 2-way ANOVA with Tukey post-hoc correction for multiple comparisons. Mice in the vehicle-treated condition are also included in Fig. 3B. Error bars represent s.e.m. Full length Western blots are shown in Supplementary Fig. 5a,c. See methods for blinding procedure.

Training-induced ERK hyperphosphorylation (Fig. 3b) coupled with increased clustered-spine stabilization (Fig. 2a) strongly implicate elevated ERK signaling in the enhanced motor learning phenotype of the *MECP2* duplication syndrome mouse model^6,34^. To further test this hypothesis, we administered a low dose (32mg/kg intraperitoneally) of the specific centrally-acting MEK inhibitor SL327 (shown in Fig. 3a) thirty minutes prior to rotarod training on each day^41^. A low dose was chosen by design to ensure effects on locomotor function in WTs are minimal if any^42^. Accordingly, while the performance of WT mice was not affected by the drug, SL327 completely reversed the enhanced motor learning in mutants (Fig. 3c, *P* = 10^−6^, genotype x drug interaction, F_genotype x treatment_ = 21.6, n=7–9 mice per group, mixed effects repeated-measures ANOVA). Furthermore, M1 ERK phosphorylation was normalized in SL327-treated mutants vs. vehicle-treated mutants (Fig. 3d, Supplementary Fig. 5c, *P* < 0.05, n=3 mice/group, 2-way ANOVA with Tukey post-hoc correction for multiple comparisons), confirming the effectiveness of the inhibitor in reducing ERK phosphorylation levels.

#### Enhanced clustered spine stabilization in *MECP2*-duplication mice is ERK dependent

Lastly, we measured the effect of MEK inhibition on learning-associated spine plasticity in *MECP2*-duplication mice (Fig. 4a). While learning-associated spine formation was not affected by SL327 (Vehicle: 4.7 ± 0.4, SL327: 4.3 ± 0.4 spines/100μm, n=6 mice per group, Fig. 4b), spine stabilization was significantly decreased (Vehicle: 1.7 ± 0.3, SL327: 1.1± 0.2 spines/100μm, Fig. 4c), back to levels similar to that of WT controls in the first experiment, reproduced from Fig. 1c-d for comparison. Separating stabilized spines into clustered and non-clustered subgroups revealed that the decrease in spine stabilization mediated by SL327 was due entirely to decreased stabilization of clustered spines (Fig. 4d). Whereas in vehicle-treated *MECP2*-duplication mice 46±3% of clustered spines were stabilized, in SL327-treated mutants only 21±3% of clustered spines were stabilized (*P*= 0.015, Mann-Whitney U test, data not shown). This observation further suggests that ERK signaling drives enhanced learning via the structural stabilization of learning-associated clustered synapses in the *MECP2*-duplication mouse.

**Figure 4.**
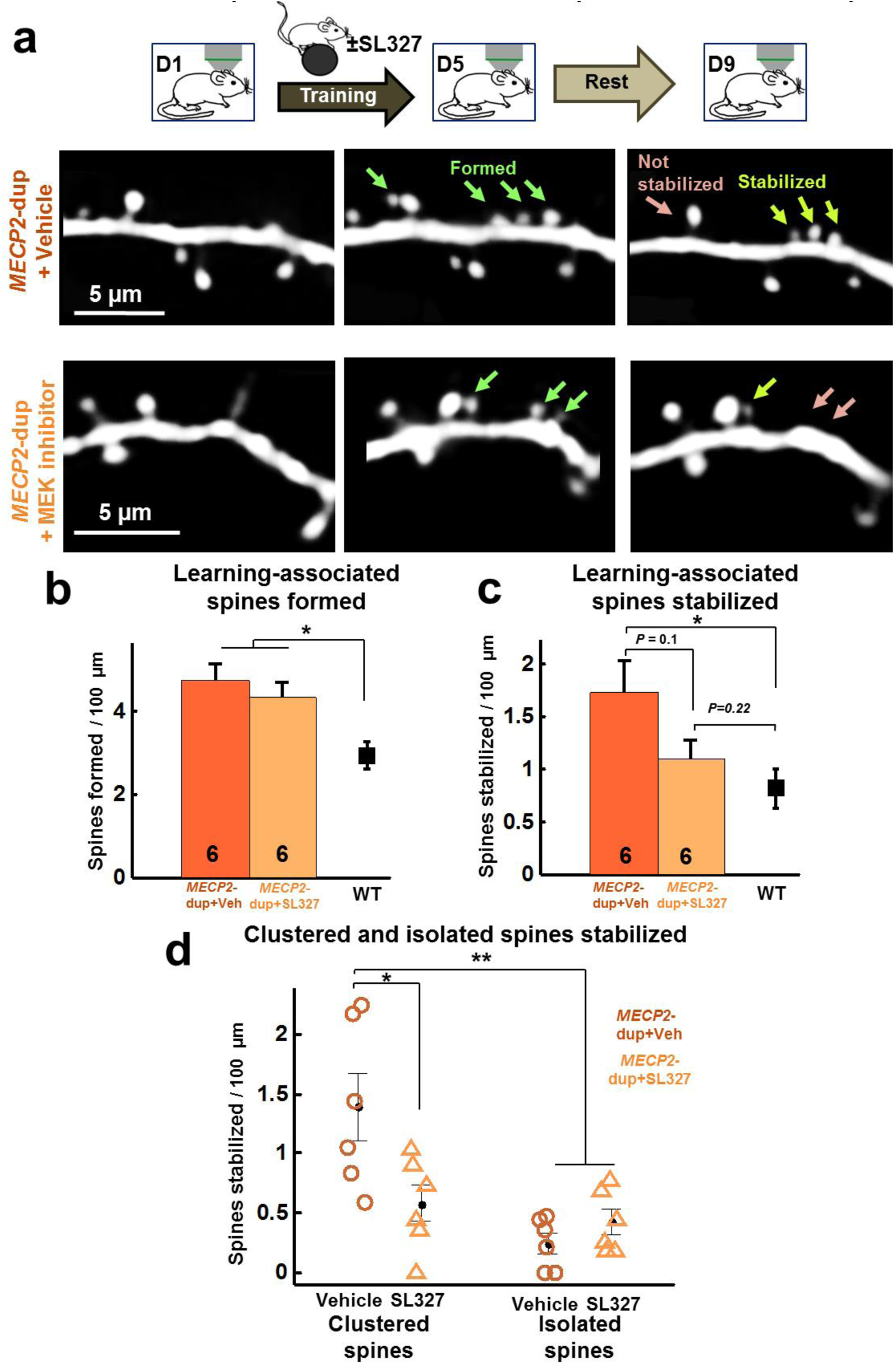
Enhanced clustered spine stabilization in *MECP2*-duplication mice is ERK dependent. **a**, Sample images of dendritic segments imaged before rotarod training (left), following 4 days of training (middle), and 4 days after the end of training (right) demonstrating decreased clustered spine stabilization in SL327-treated (bottom) vs. Vehicle treated (top) *MECP2*-duplication mice. **b**, Learning-associated spines formed per 100μm in vehicle-treated (dark orange, n=6 animals, 184 spines formed, 54 dendritic segments) and SL327-treated (light orange, n=6 animals, 143 spines formed, 50 dendritic segments) *MECP2*-duplication mice. Original data from WT animals (black) is included for comparison. * *P* < 0.05, F(2,19)=7, 1-way ANOVA or Mann-Whitney U test. **c**, Learning-associated spines stabilized per 100μm in each condition. * *P* < 0.05 F(3,8)=4.56, oneway ANOVA or Mann-Whitney U test. d, Number of clustered and isolated spines (see Methods) stabilized per 100μm in vehicle-treated versus SL327-treated *MECP2*- duplication mice. Data points are from individual animals. * *P* = 0.018, ** *P* < 0.01, F_clustering_(1,20)=13.7, F_treatment_(1,20)3.27, F_clustering x treatment_(1,20)=7.9, one-way ANOVA across animals with Tukey test for multiple comparisons. Note that there are significantly fewer learning-associated stabilized clustered spines in SL-327 treated animals.

### Discussion

In summary, we found that the abnormal repetitive motor learning phenotype in *MECP2*-duplication mice is associated with excessive cooperative stabilization of new synaptic connections that form in clusters in primary motor cortex during learning. This suggested a dysregulation in cell signaling pathways that drive cooperative dendritic spine stabilization in clusters. The ERK pathway, a major regulator of clustered spine stabilization^34^, was indeed hyperactive in *MECP2*-duplication mouse motor cortex specifically after motor training. Furthermore, both increased spine-cluster stabilization and enhanced motor learning in *MECP2*-duplication mice depend on hyperactivity of the ERK pathway and can be reversed by ERK-specific pharmacologic inhibition.

#### Tg1 mice show enhanced cooperative learning-associated dendritic spine stabilization

Enhanced repetitive motor learning on the rotarod is observed in several autism mouse models with prominent behavioral inflexibility, including the *MECP2*- duplication^6, 8^, neuroligin-3^13^, 15q duplication^14^, PTEN^15^, and CNTNAP2 mice^43^, providing a robust model behavior for studying the abnormal consolidation of repetitive motor routines. These mouse models also demonstrate higher spontaneous dendritic spine turnover rates^17–20^, suggesting they share a deficit in the balance between structural synaptic plasticity and stability. How this balance is affected by learning has, so far, not been studied in these models. Our goal is to forge a link between abnormal structural synaptic plasticity and the behavioral phenotype of enhanced repetitive motor learning seen in the *MECP2*-duplication model of autism. We focused on studying corticospinal neurons in the primary motor cortex, the final common motor pathway that integrates learning effects throughout the motor circuit to promote optimal movement control^44^.

New dendritic spines are known to form on the apical tufts of corticospinal neurons in area M1 of WT mice during motor learning^21,22^. A fraction of these spines get stabilized over time, while others disappear. The number of stabilized spines is known to correlate with motor performance in healthy wild type mice. Therefore the new spines stabilized with motor training are thought to represent a structural correlate of procedural motor learning.

This paradigm allowed us to investigate whether or not the formation and stabilization of new dendritic spines associated with procedural learning is abnormal in autism. We found that ~33% more new spines are formed in apical tufts of *MECP2*-duplication mouse corticospinal neurons during motor learning compared to littermate controls. These learning-associated spines are ~40% more likely to be stabilized compared to controls, leading to almost twice as many new spines stabilized with learning in *MECP2*-duplication animals.

In WT mice, spines formed in area M1 during motor learning aggregate in clusters on corticospinal neuron dendrites, arguing that synaptic clusters constitute a subcellular substrate for procedural learning^33^. Interestingly, we found that approximately 3 times as many learning-associated spines are stabilized within 9 □m long clusters in *MECP2*- duplication mice compared to littermate controls. Furthermore, stabilization of clustered <u>*but not of isolated*</u> spines correlates well with the *MECP2*-duplication mouse’s enhanced rotarod performance, suggesting that an increase in cooperative spine stabilization underlies this mutant’s abnormal motor learning phenotype.

An increase in clustered spine stabilization can potentially have important functional implications^32^. Clusters of synapses drive neuronal activity more strongly when activated synchronously through nonlinear dendritic integration mechanisms^45^. Neurons that implement synaptic clustering may fire selectively to precise combinations of inputs spatially co-localized on the dendrite, in theory dramatically increasing memory storage capacity^46^. Too much input clustering, however, may potentially lead to “overfitting” of learned representations leading to a rigid and restricted behavioral repertoire that may not be flexible enough to accommodate the efficient learning of new representations.

It is interesting to speculate on the origins of the presynaptic inputs to the learning-associated corticospinal apical-tuft dendritic spines we studied. Corticospinal neurons integrate information from premotor cortex, somatosensory cortex, and corticostriatal and corticocerebellar circuits to implement adaptive motor control^47–49^. Abnormal plasticity in these networks could therefore contribute to enhanced behavioral learning that may manifest on corticospinal neuron dendrites as modified synaptic connectivity. In this context, we note the recent report that enhanced motor learning in the neuroligin-3 mutant mouse model of autism arises from hyper-excitability in direct pathway medium spiny neurons of the ventral striatum^13^. It is intriguing that *MECP2* appears to suppress neuroligin-3 expression in vivo^50^ and ventral striatum connects to motor cortex via thalamically-relayed projections that could be presynaptic to corticospinal neurons in layer 1^51^. However, several pre-synaptic pathways that project to corticospinal neurons in L1 could in principle contribute to our observations. It is of considerable interest to dissect the individual contributions of these pathways specifically, in the future.

Abnormal learning and synaptic plasticity are highly penetrant features of autism mouse models. Our data provides the first in vivo evidence that: 1) abnormal learning-associated synaptic consolidation contributes to the altered motor learning phenotype of the *MECP2*-duplication mouse model of autism, and 2) cooperative spine stabilization leading to the enhanced formation of learning-associated synaptic clusters correlates strongly with the level of behavioral performance. Identifying the molecular mechanisms that underlie these abnormalities in synaptic plasticity may lead to candidate therapeutic targets in the future.

#### ERK signaling mediates the observed increase in synaptic clustering

The Ras-MAPK (Ras-Raf-MEK-ERK) pathway is a ubiquitous signaling pathway by which extracellular stimuli modulate synaptic plasticity and cellular physiology^37^. Ras-MAPK genes are dysregulated in *MECP2*-duplication mice^39^, mutations in Ras-MAPK pathway genes are linked to several forms of autism^38^, and several autism models have been shown to have abnormal Ras-MAPK signaling^52^.

Ras-MAPK signaling in hippocampal pyramidal neurons has been shown in vitro to be specifically involved in the cooperative potentiation of neighboring dendritic spines^34,53,54^. When a spine is sufficiently activated, Ras enters its GTP-bound active state and diffuses 9–10 μm down the dendrite to invade neighboring spines. There it initiates the ERK phosphorylation cascade (Fig. 3a) orchestrating a number of transcriptional and translational changes associated with synaptic consolidation^37^.

Our findings of 1) increased cooperative stabilization of learning-associated spines in 9–10 μm clusters whose number correlates with enhanced motor learning, 2) increased M1 ERK signaling specifically after motor training, 3) reversal of enhanced motor learning by specific ERK inhibition, and 4) normalization of learning-associated clustered spine stabilization by ERK inhibition, provide a compelling basis for the idea that elevated ERK drives enhanced motor learning in *MECP2*-duplication mice through the stabilization of learning-associated dendritic spine clusters.

These results traverse genetic, cell signaling, microcircuit, and behavioral levels of pathophysiology to link an autism-associated mutation to an altered learning phenotype. Moreover our results provide the first *in vivo* evidence that ERK signaling is involved in learning-associated synaptic clustering^33,36^ and suggest that a training-dependent increase in M1 ERK signaling facilitates procedural motor memory consolidation.

In future work it will be fruitful to study the regulation of upstream mediators of ERK signaling^37,39,52^, to begin to understand how it is that ERK becomes dysregulated in *MECP2*-duplication mice after training^55,56^.

#### Two classes of autism mouse models defined by learning and spine clustering

Opposite to our findings in the *MECP2*-duplication mice, it was recently reported that Fragile X mice have *impaired* motor learning and *decreased* clustered spine stabilization^57–59^ We propose that the divergent learning and plasticity phenotypes of these mouse models typify two distinct classes of syndromic autism (Supplementary Fig. 6). The first group, exemplified by *MECP2*-duplication, neuroligin-3 mutants, and 15q duplication mice, exhibits enhanced motor learning^6,13,14^, prominent behavioral inflexibility, increased learning-associated clustered spine stabilization (shown so far only in *MECP2*-duplication mice), enhanced LTP^6,60,61^, and increased activity-dependent ERK signaling (also shown so far only in *MECP2*-duplication mice). The second group, typified by Fragile X, Rett syndrome, and 15q deletion (Angelman) mice, demonstrate impaired motor learning^57,62–64^, decreased learning-associated clustered spine stabilization (shown only in Fragile X mice)^59^, impaired LTP^64–66^, and decreased activity-dependent Ras-MAPK signaling^67^. This proposed taxonomy expands upon the previous demonstration that Fragile X syndrome and tuberous sclerosis form an axis of synaptic pathophysiology^68^, and makes several experimental predictions, summarized in Supplementary Fig. 6.

#### Implications for autism pathophysiology

Changes in synaptic stabilization that have positive effects early in the *MECP2*-duplication mouse’s postnatal development likely become detrimental later in life, producing spasticity, behavioral inflexibility, and epilepsy^6^. Although enhanced motor learning is not observed in human patients^11^, it is possible that the structural abnormality we describe in mice is also operating in the humans, but manifests with a different developmental trajectory of motor deterioration due to species-specific differences in motor system development and function^69^. Our finding that an inhibitor of Ras-MAPK signaling was able to normalize both clustered spine stabilization and motor learning in *MECP2*-duplication mice suggests that modulation of structural plasticity through Ras-MAPK inhibitors may provide a potential therapeutic avenue for some manifestations of the *MECP2* duplication syndrome.

These findings exemplify how neural circuit analysis can generate hypotheses that implicate specific molecular mechanisms of disease. Our results lend support to recent hypotheses implicating the ERK signaling pathway in autism pathophysiology^38,52,70^, and suggest that increased clustered-spine stability may contribute to some of the symptoms in *MECP2* duplication syndrome. This provides evidence that the exceptional learning capabilities seen, at times, in autism^2,13,14^ may arise as a result of increased learning-associated synaptic stability. In the future, it will be valuable to explore further the proposition that a pathological imbalance between synaptic stability and plasticity in different circuits might account for different phenotypic aspects of autism spectrum disorders.

### ONLINE METHODS

#### Animals

FVB-background *MECP2* duplication (Tg1) mice ^6^, were crossed to C57 thy1-GFP-M^26^ homozygotes obtained from Jackson Laboratories, to generate F1C57;FVB MECP2-duplication;thy1-GFP-M mice and thy1-GFP-M littermate controls. Males were used in experiments reported in Fig. 1–3; Data reported in Fig. 4 were acquired from 4 males and 2 females per condition. Animals were housed in a 12/12 light/dark cycle (lights on from 7 am to 7 pm). All experiments with animals were carried out in accordance with the National Institutes of Health guidelines for the care and use of laboratory animals and were approved by the Institutional Animal Care and Use Committee of Baylor College of Medicine.

#### Blinding

In data reported in Fig. 1–2, surgeries, imaging, rotarod training, and analyses were performed blind to genotype. Mouse numbers and genotypes were placed in a 2-column spreadsheet by a lab member not involved in the experiment or analysis (J. Park), and Matlab scripts imported genotypes from this spreadsheet to plot data without unblinding the experimenter (R.T.A.) to genotypes. In data reported in Fig. 3b,d, mice from each genotype were randomly assigned to the drug or vehicle condition. Material to be injected was placed into individual tubes for each animal, then mice were injected without knowledge of the test tube’s contents. Mice were trained, motor cortex was harvested, and Westerns were run and analyzed blind to condition. In Fig. 4 data, only *MECP2*-duplication mice were used, and imaging, rotarod training, and drug/vehicle injection was performed unblinded to condition. Prior to analysis however, data sets from each animal were assigned a new blinded number by a lab member not involved in the experiment or analysis (J. Park), and spine counting and analysis were performed blind to condition.

#### In vivo two-photon imaging

Surgeries and imaging were performed blind to genotype. At least two weeks prior to the first imaging session (~12–14 week-old-mice), a 3 mm-wide opening was drilled over motor cortex, centered at 1.6 mm lateral to bregma based on ^28^, and a glass coverslip was placed over the exposed brain surface to allow chronic imaging of neuronal morphology ^24,71^. Dendritic spines were imaged using a Zeiss in vivo 2-photon microscope with Zeiss 20x 1.0 NA water-immersion objective lens. High-quality craniotomies had a characteristic bright-field appearance with well-defined vasculature and pale grey matter. Under two-photon scanning fluorescent dendrites were reliably clear and visible with low laser power (<20 mW on the pia) and photomultiplier tube voltage.

Only high quality preparations (low background noise across all time points, <5 pixel [0.25 μm] motion artifact, and dendrites well isolated from other fluorescent structures) were used in the blinded analysis (Supplementary Fig. 1a, Supplementary Movie 1). GFP-labeled neurons were first imaged at low resolution (620×620 μm FOV, 0.6 μm/pixel in XY, 2.5 μm Z-step size) down to 600–700 μm to capture all of the cell’s dendritic processes and assay cell subtype by morphology, primary apical bifurcation depth, and soma depth ^72^ The apical dendrites from complex-tufted neurons, the corticospinal neurons projecting to the spinal cord and thalamus in M1 ^27,73^, were selected based on their large highly ramified dendritic trees, deep primary apical bifurcation, and thick dendrites ^72^, and re-imaged at high resolution (310×310 to 420×420 μm FOV, 0.1 μm/pixel, 1 μm Z-step size) to adequately capture individual dendritic spines. Laser power was maintained under 20 mW (average ~10 mW) during image stack acquisition.

#### Analysis of structural plasticity

Raw **z** stacks were denoised by a custom polynomial interpolation method^17^. Learning-associated spine formation, elimination, and stabilization were quantified with a custom Matlab user interface and ImageJ (MicroBrightField). Terminal dendrite segments which were well-visualized at all time points were chosen for analysis. Sections of dendrite occluded by other fluorescent structures or blood vessels were excluded from the analysis. Due to in vivo two-photon microscopy’s relatively poor resolving power in the z-axis, only structures protruding laterally along the X-Y plane were included in the analysis, following the standard in this field ^24^ For a protrusion to be selected for analysis it had to project out of the dendritic shaft by at least 4 pixels (~0.4 microns), which corresponds approximately to 2 SDs of the noise blur on either side of the dendritic shaft. Spines were initially identified at one time point, by moving up and down through individual slices in each Z stack, and the same region of dendrite was examined at other time points to identify the first (formation) and last (elimination) time that the spine was present. Custom Matlab routines analyzed the stability/survival of each formed spine. Filopodia, which are rare at the analyzed age, were identified morphologically, based on their long length (usually >4 μm), curved shape, and lack of a distinct head ^74^ and excluded from the analysis as in ^21^.

#### Analysis of dendritic spine clustering

Each spine was classified as either clustered or isolated by calculating the distance to its nearest-neighbor stabilized learning-associated spine ^33^ The difference in clustered spine stabilization rate between *MECP2*-duplication and WT mice was calculated as a function of cluster distance threshold (max distance to nearest co-stabilized spine to be counted as clustered) (Supplementary Fig. 4). This difference in clustered spine stabilization increased with increasing cluster distance threshold and plateaued at 9 μm; we used this threshold to define spine clusters for further analysis ^34,35^.

#### Spine clustering simulations

Simulations of one thousand dendrites per genotype were generated based on the measured mean and standard deviation of dendritic length, the number of baseline spines and formed spines, and the spine stabilization rate in each genotype. Spine locations were simulated in one dimension. The same nearest neighbor clustering analysis was performed on these simulated dendritic spines as was performed on the actual data. Results are shown in Supplementary Fig. 3a.

#### Motor training

The Ugo Basile mouse rotarod was used for motor training. At least two hours after imaging sessions, in the late afternoon, mice were placed on the rotarod, and the rotarod gradually accelerated from 5 to 80 rpm over 3 minutes. Single-trial rotarod performance was quantified as the time right before falling or holding on to the dowel rod for two complete rotations without regaining footing. A 5–10 minute rest period occurred between each trial. Four trials were performed per day. Per-animal rotarod performances used in scatter plots (Fig. 1f, 2c,d) were calculated as the best median per-day performance. For immunoblot experiments measuring M1 ERK phosphorylation with rotarod training, mice were trained 8 times (Fig. 3b) or 6–8 times (Fig. 3b,d). For the pharmacology experiments demonstrating the effect of the specific MEK inhibitor SL327 on rotarod performance, the peak performance of each animal on each day, normalized to the vehicle-treated WT mean, was used in the analysis (median performance yielded similar statistically significant results).

#### Pharmacological Ras-MAPK inhibition

The centrally-acting selective MEK inhibitor SL327 (Axonmedchem #1122) ^75^ was injected intraperitoneally at 32 mg/kg (in 16 mg/mL DMSO) 30 minutes prior to rotarod training ^41^. This dose was selected to be low as it is known to minimally affect motor performance in wild-type animals ^41^.

#### Microstimulation mapping of motor cortex

After the final imaging session, microstimulation mapping of motor cortex was performed (see Supplementary Fig. 1b) as described in ^28^ Under ketamine/xylazine anesthesia, the coverslip was removed from the craniotomy, and a bipolar stimulating electrode was lowered into the imaged brain region in a 0.5×0.5 mm grid pattern to ~750 μm depth (deep L5). Current pulses starting at 10 μA (up to 60 μA) were applied until twitching was observed in contralateral muscle groups. Twitching muscle groups were scored as forelimb, hindlimb, face, tongue, or tail. Imaged dendritic arbors were registered with mapped regions using blood vessels as landmarks. All structural plasticity data were acquired from forelimb and hindlimb motor cortex.

#### Immunoblots

Deeply anesthetized (isoflurane) 4–5 month old *MECP2*-duplication mice and littermate controls were sacrificed 30 minutes after training and their brains were rapidly dissected on a glass plate over ice. The motor cortex from each hemisphere was isolated from the remaining cortical tissue and lysed in ice-cold homogenization buffer [in mM: 200 HEPES, 50 NaCl, 10% glycerol, 1% Triton X-100, 1 EDTA, 50 NaF, 2 Na_3_VO_4_, 25 β-glycerophosphate, and 1x EDTA-free complete ULTRA protease inhibitor cocktail tablets (Roche, Indianapolis, IN)]. Insoluble material was removed by centrifugation at 14,000*g* for 10 minutes at 4°C. Protein concentration of the resulting supernatant was determined by Bradford assay (Biorad, reagent 500-0006) and lysates were then diluted in 2x Laemmli Buffer. A total of 30 μg protein/sample was resolved by SDS-PAGE (12.5% acrylamide) and gel contents were transferred to nitrocellulose membranes. Membranes were blocked 30 minutes in 5% milk, 0.2% Tween-20 tris-buffered saline (TBST). To assess ERK phosphorylation, membranes were first probed overnight at 4°C with phospho-specific rabbit anti-phospho-p44/42 MAPK/Erk1/2 (Cell Signaling Technology, #4370, 1:1000). Blots were then incubated with HRP-conjugated goat anti-rabbit secondary antibody (Jackson ImmunoResearch, 111-035-144, 1:5,000) for 1h at room temperature followed by incubation in Super-Signal West Femto kit substrate (Thermo Scientific, 34096) per the manufacturer’s instructions. Film was exposed to the Super-Signal-treated membranes and then developed. Membranes were then stripped (1.5% glycine, 2.9% NaCl, pH2.8), blocked with 5% milk TBST, then re-probed overnight with rabbit anti-MAPK/ERK1/2 (Cell Signaling Technologies, #9202, 1:1000) to assess total ERK protein levels. β-actin levels were measured as an additional loading control (mouse anti-β-actin, Millipore, MAB1501, 1:5000). The goat anti-mouse secondary antibody was diluted 1:10,000 for the β-actin blots. Band density was quantified in ImageJ (NIH).

#### Image presentation

Dendritic spine images are displayed as ‘best’ projection mosaics. Extraneous fluorescence is masked and images are slightly smoothed for illustration purposes only.

#### Statistical tests

Statistical significance between unpaired normal samples was assessed by two-tailed student’s t-test, and with Mann-Whitney U test for non-normal samples, except where noted. Normality was assessed by the Kolmogorov-Smirnov test. Enhanced rotarod performance in *MECP2*-duplication mice in Fig. 1 was tested by repeated-measures ANOVA. Effect of SL327 on rotarod performance in *MECP2*-duplication mice compared to littermate controls was evaluated by determining the genotype x treatment interaction term using a mixed effects repeated-measures ANOVA in R statistics software. Statistical analysis was performed on raw per-trial rotarod performance values. Litter was included as a term to control for across-litter variability in performance. Statistical significance of the correlation between rotarod performance and spine stabilization was assessed using the Pearson correlation coefficient and student’s t-distribution ^76^ \**P*≤0.05, \*\**P*≤0.01, \*\*\**P*≤0.001. All results are reported as mean ± sem, unless otherwise noted.

## AUTHOR CONTRIBUTIONS

R.T.A., S.M.S., and S.A.B. designed the experiments and wrote the manuscript. R.T.A. conducted the in vivo imaging and rotarod training experiments and analyzed the spine stabilization and rotarod performance data; S.A.B conducted and analyzed the immunoblotting experiments. H.Y.Z. contributed to the design of the project, review and critical discussion of data, and editing the manuscript. M.C.M. contributed to the design of the SL327 MEK inhibitor experiments.

## ACKNOWLEDGMENTS

We are grateful to J. Park, G. Allen, S. Torsky, E. Sztainberg, B. Suter, J. Meyer, A. Palagina, J. Patterson, S. Shen, D. Yu, and K. Tolias for technical and theoretical advice on experiments and comments on the manuscript. We thank H. Lu for the images of mouse coronal brain slices.

R.T.A. received support from the Autism Speaks Weatherstone Fellowship and the BCM Medical Scientist Training Program. This work was supported by grants from the Simons Foundation (240069, SS) and March of Dimes to S.M.S., the Howard Hughes Medical Institute and NINDS HD053862 to H.Y.Z., and the Baylor Intellectual and Developmental Disabilities Research Center (P30HD024064) Mouse Neurobehavioral Core.

## SUPPLEMENTAL FIGURES AND LEGENDS

**Supplementary Figure 1.**
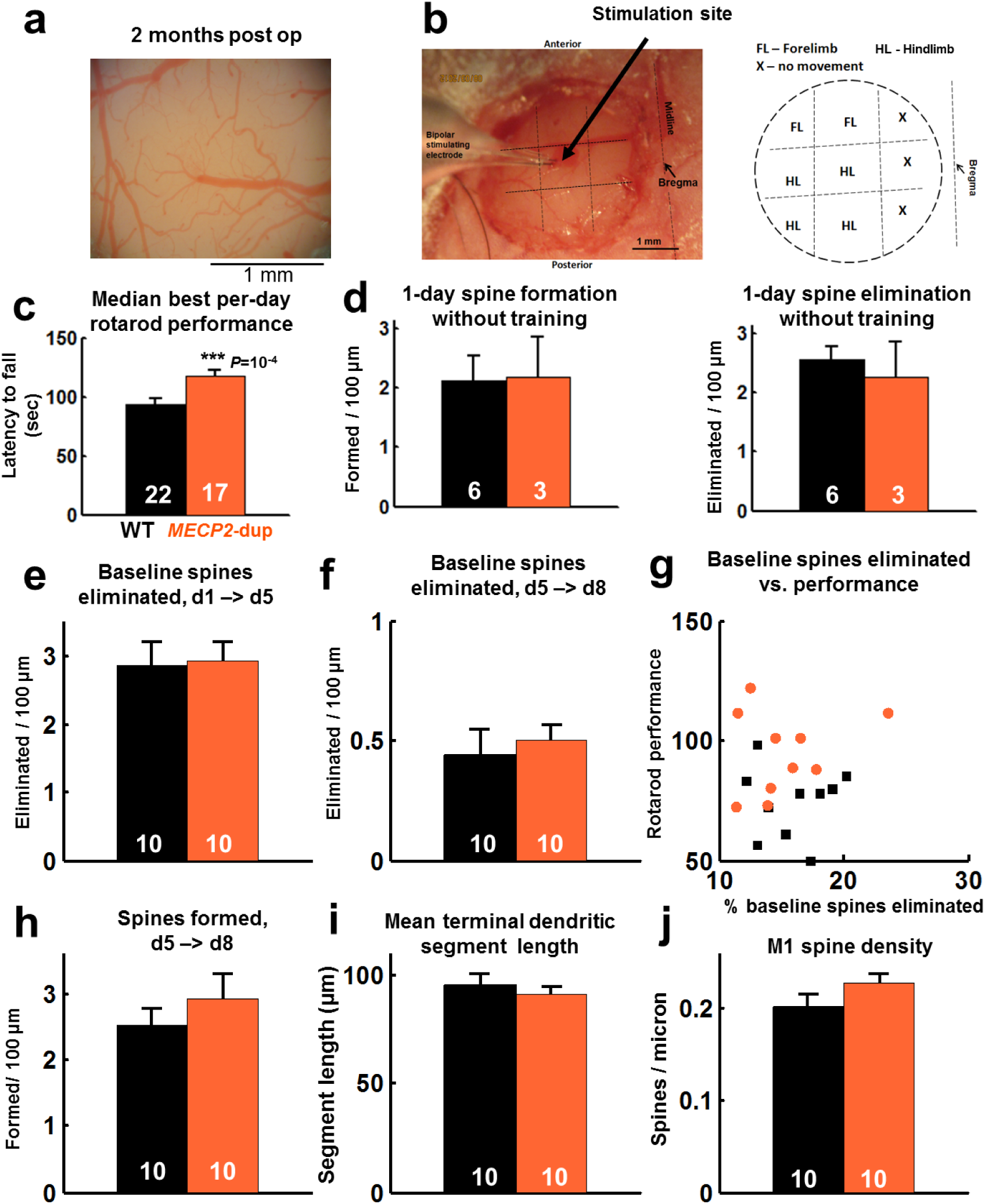
Supplemental Structural Plasticity Graphs, Related to Fig. 1. **a,** Example brightfield image of chronic cranial windows over M1 at 2 months post-op, showing the well-defined vasculature and pale grey matter characteristic of high-quality preparations. **b,** *Left,* image demonstrating the microstimulation experiment, performed post-hoc in experimental mice at the end of imaging. A bipolar stimulating electrode was lowered ~750 μm into the window at 9 sites in a 1000 micron grid. *Right,* an example map of motor cortex generated by microstimulation. Forepaw and hindpaw twitches were generated at low currents in all stimulated cortices (n=10 mice), confirming localization to M1. **c,** Median best per-day rotarod performance in mutants (orange, n=17) and WT (black, n=22). Error bars represent mean±sem. P=10^−4^, t test. **d,** *Left,* Spontaneous 1-day spine formation rate in *MECP2*-duplication mice (orange) versus WT controls (black). *Right,* Spontaneous 1-day spine elimination rate. Note the difference without training did not reach significance in the motor cortex at this age group. **e,** Baseline (pretraining) spines eliminated after 4 days training and 4 days rest. **f,** Baseline spines eliminated in the 4 days that follow the period of rotarod training. **g,** Percent of baseline spines eliminated versus rotarod performance per animal. **h,** New spines formed in the 4 days that followed rotarod training **i,** Histogram of terminal apical tuft dendritic branch lengths in M1. **j,** Spine density of analyzed terminal dendritic branches in the apical tuft of corticospinal neurons in area M1. P values calculated with Mann-Whitney U test. None of these comparisons from c to h reached statistical significance. Note that these results differ in some respects from previous measurements in the somatosensory cortex of *MECP2*-duplication mice ^17^, potentially due to differences in brain region, dendrite selection criteria, or imaging duration (1 day in this study vs. 1 hour in the previous study).

**Supplementary Figure 2.**
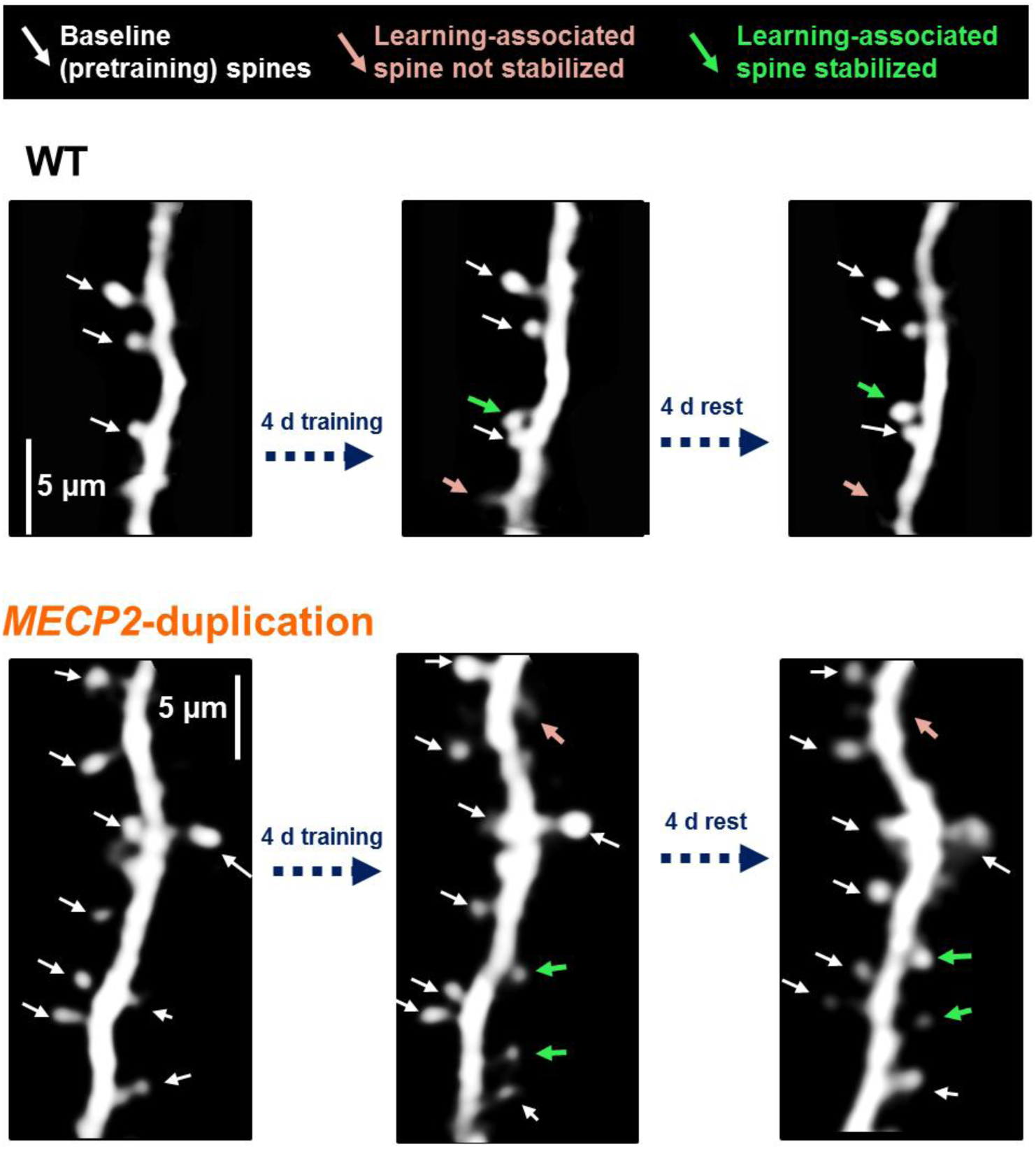
Following Dendritic Segments Across training, Related to Fig. 1. Example dendritic segments at baseline prior to training (left), four days after rotarod training (middle), and four days after the end of training (right). Baseline spines prior to training are denoted by white arrows, learning-associated spines that do not stabilize by pink arrows, and learning-associated spines that do stabilize by green arrows. *Top:* WT control. *Bottom: MECP2*-duplication mouse. Stabilized spines cluster along the dendrite in mutants (See also population results in Fig. 2a). Scale bars: 5 μm.

**Supplementary Figure 3.**
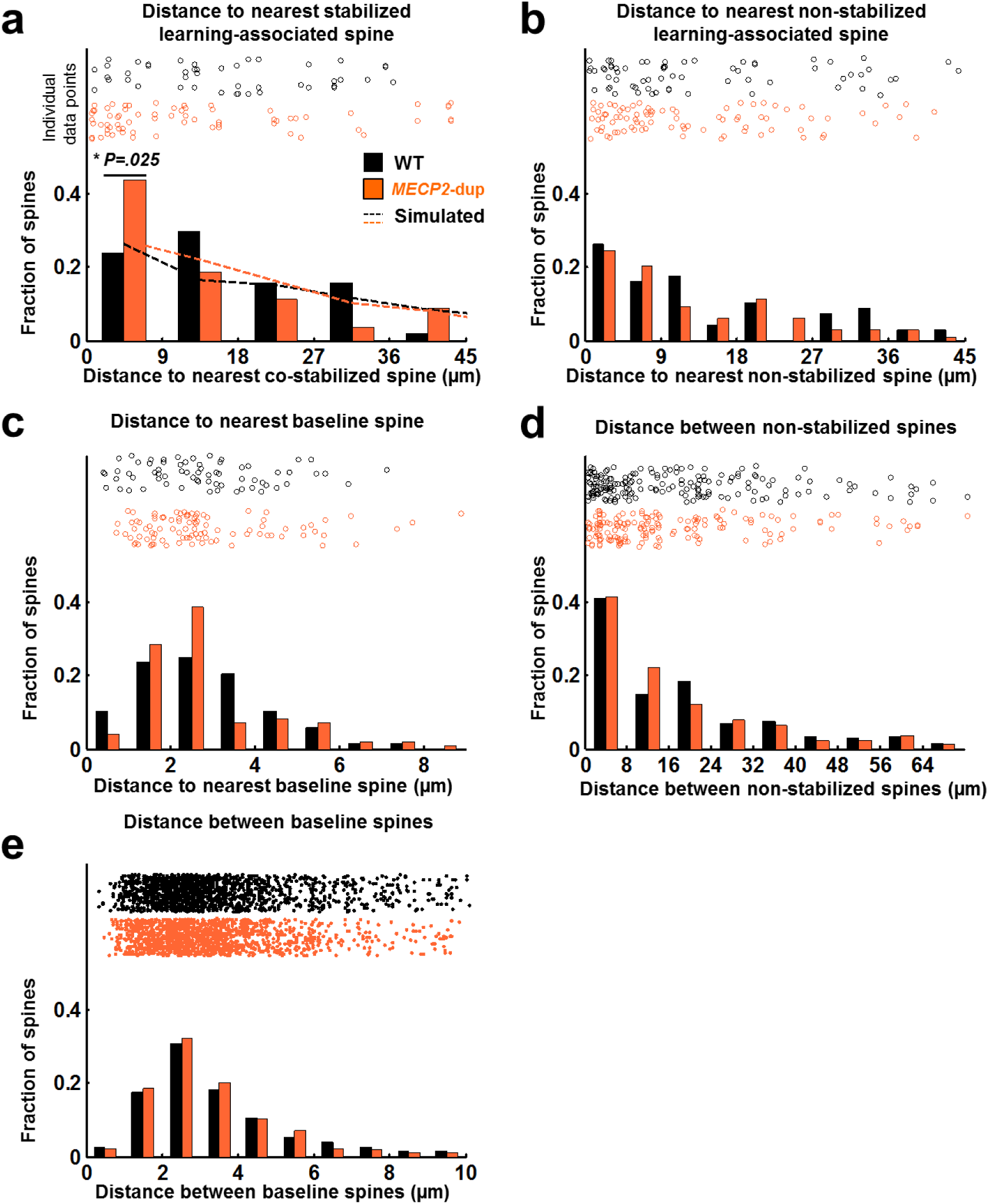
Supplemental Clustered Synaptic Plasticity Graphs, Related to Fig. 2. **a,** Histogram of the distance to the nearest co-stabilized spine for each stabilized learning-associated spine, in *MECP2*-duplication (orange) and WT (black) mice. Individual data points are shown at the top. Twice as many spines were co-stabilized within nine microns of each other overall in mutants. * *P*=0.025, Fisher exact test. Dotted lines depict estimated distances between stabilized spines simulated from the number of spines formed per micron in each genotype (see Methods). Differences in overall spine formation and stabilization in mutants do not explain the increase in clustered spine stabilization between mutant and WT (*P*=0.65, Fisher exact test). **b,** Histogram of distances between each learning-associated stabilized spine to the nearest *<u>non-stabilized</u>* learning-associated spine. c, Histogram of distances from each learning-associated stabilized spine to the nearest pre-existing baseline spine. d, Histogram of nearest-neighbor distances between *<u>non-stabilized</u>* learning-associated spines. e, Histogram of nearest-neighbor distances between baseline (prior to training) spines. None of the distributions in b-e showed significant differences. WT: n=76 learning-associated spines, 99 dendrites, 9409 μm total dendritic length. *MECP2*-duplication: n=109 learning-associated spines, 82 dendrites, 7390 μm total dendritic length.

**Supplementary Figure 4.**
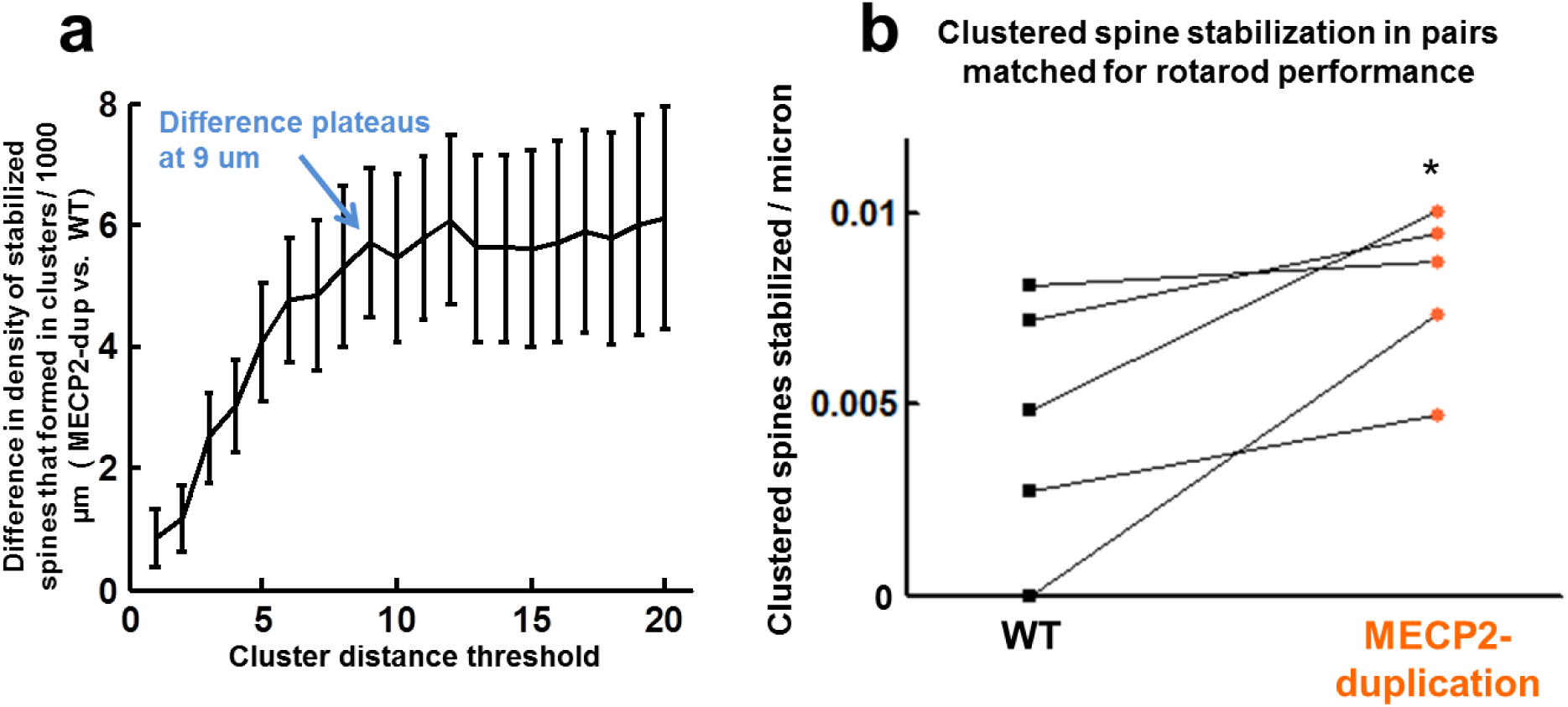
Determination of Cluster Distance Threshold, Related to Fig. 2. **a,** Increase in clustered spine stabilization in mutants vs. WT littermates, plotted as a function of the cluster distance threshold applied (i.e. the max distance to a co-stabilized learning-associated spine to be categorized as clustered). Error bars represent the summed s.e.m. from both genotypes. The difference in clustered spine stabilization between genotypes increases with increasing cluster distance threshold leveling off to a plateau at ~9 μm. 9 μm was therefore chosen for further analysis, and this agrees with the range of spine consolidation cooperativity shown in vitro by (Harvey et al., 2007). **b,** Clustered spine stabilization in pairs of WT and *MECP2*-duplication mice matched for time spent on the rotarod (<3 seconds difference in mean time before falling). Pairs are denoted by lines. Increased clustered spine stabilization is observed in Tg1 mice even when matching for time spent on the rotarod. *P* < 0.05, paired t test, n=5 per genotype. Note that only the 5 worst performing mutants and 5 best performing WT mice were able to be included in this analysis.

**Supplementary Figure 5.**
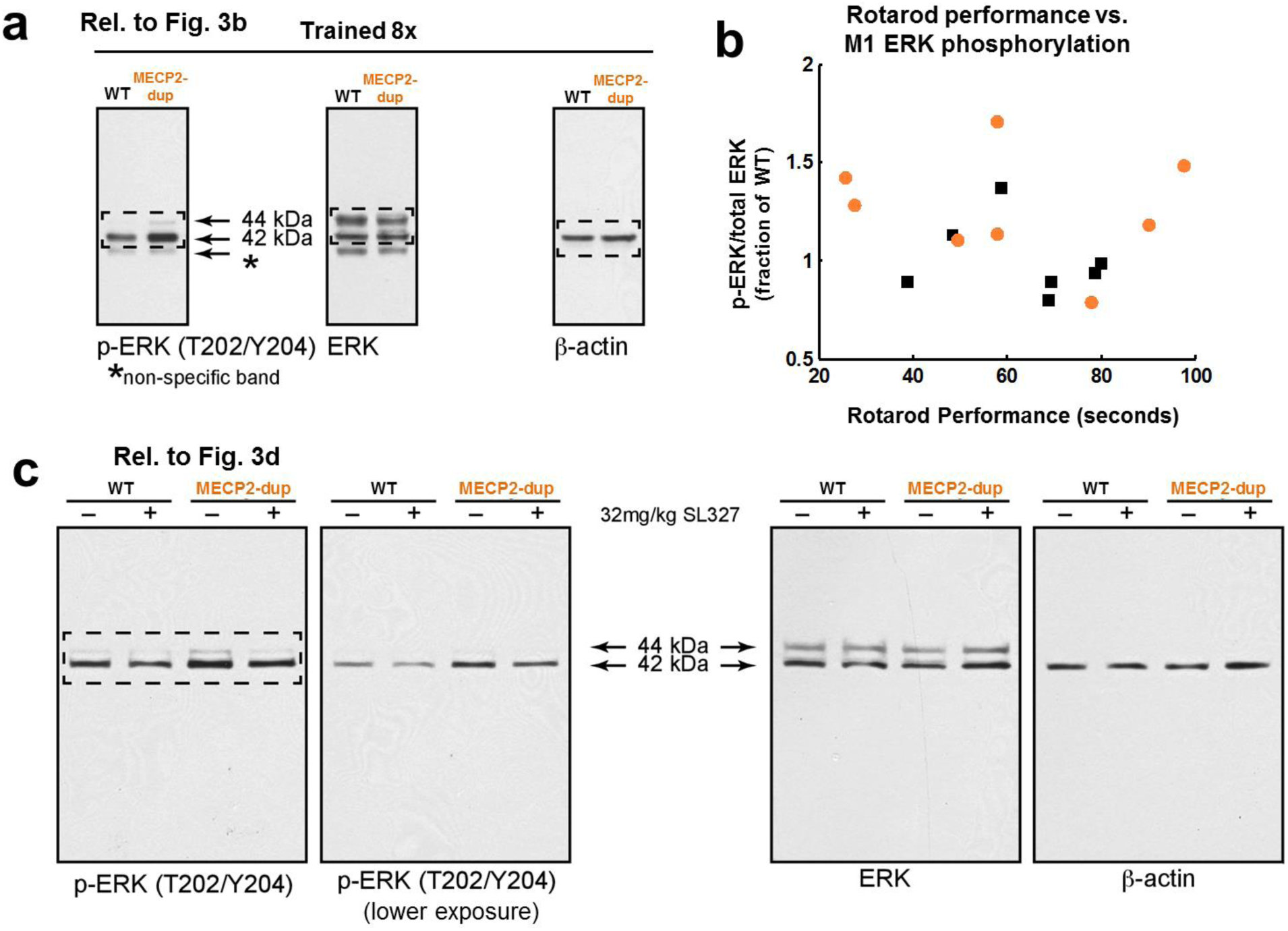
Western blot data, Related to Fig. 3. **a,** Full-length Western blots relevant to Fig 3b, showing example immunoblots to p-ERK (T202/204), total ERK, and beta-actin. **b,** Scatter plot of rotarod performance and ERK phosphorylation per animal in mutants (orange, n=8) and WT (black, n=7). ERK phosphorylation did not correlate will with rotarod performance in individual animals (r=− 0.17, p=0.53). a, Full-length Western blots relevant to Fig 3d, showing example immunoblots to p-ERK (T202/204), total ERK, and beta-actin. Note SL-327 suppresses the level of p-ERK in mutant animals.

**Supplementary Figure 6.**
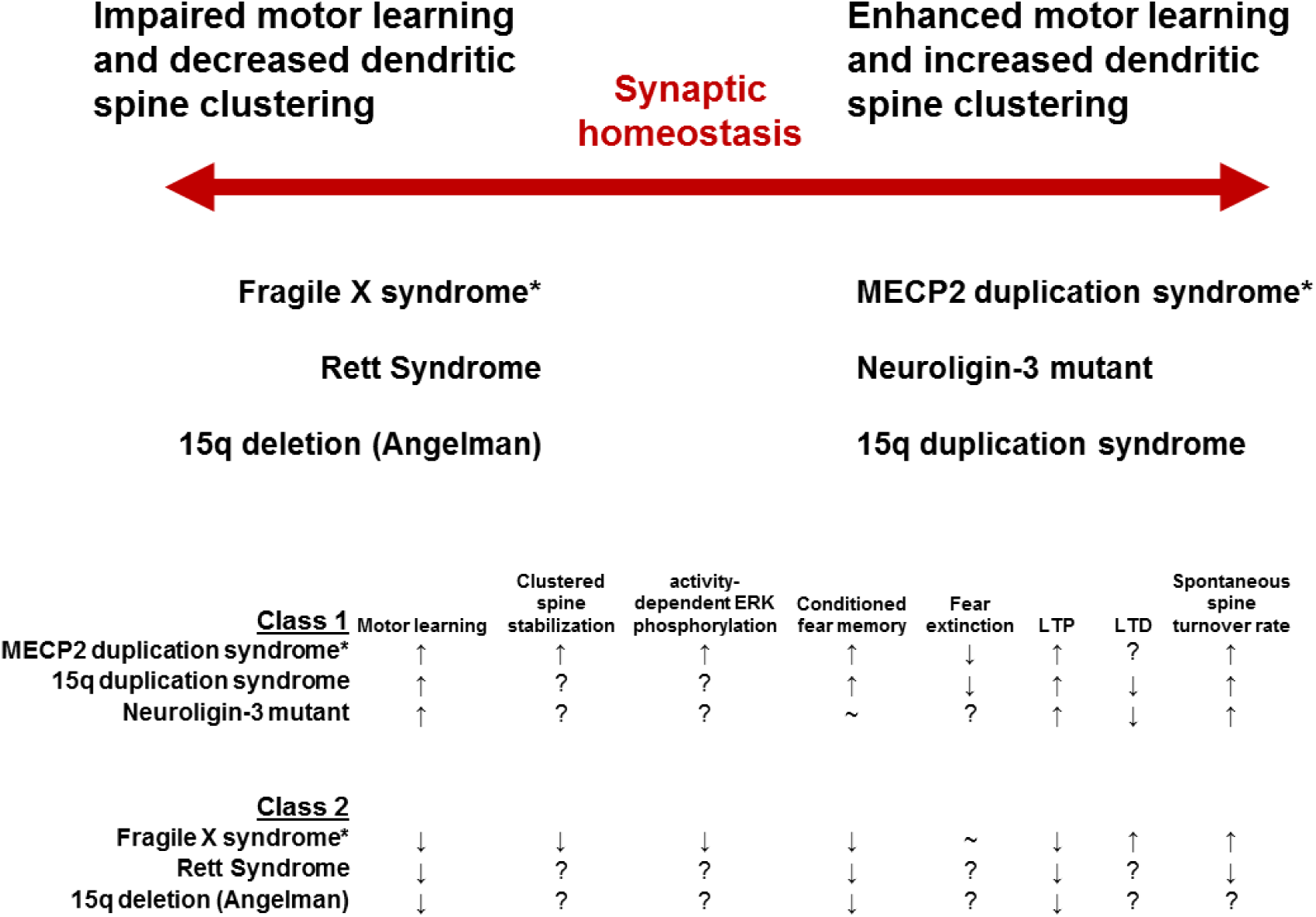
Two classes of autism models defined by learning and spine clustering. Top: Proposed classification of autism mouse models, defined by motor learning and spine clustering. Asterisks denote exemplars for each group where differences in clustered spine stabilization have been observed. Bottom: Table showing range of phenotypes shared within each group and opposite across the groups. <u>***Citations for Class 1:***</u> motor learning: ^6,13,14^, learning-associated clustered spine stabilization (Fig. 2), conditioned fear ^17,77^, fear extinction (^8^, Unpublished observations), LTP^6,60,61^, LTD: ^61,78^, activity-dependent Ras-MAPK signaling (Fig. 3). Spine turnover ^17–20^ Citations for Class 2: motor learning,^57,58^,^62–64^,^79^, learning-associated clustered spine stabilization ^59^, conditioned fear ^79,80^, LTP ^64–66^ activity-dependent Ras-MAPK signaling ^67^, spine turnover ^57,81–83^.

**Supplementary Movie 1. Example Z Stack of Raw Data Acquired by In Vivo Structural Imaging, Demonstrating Characteristic Sparse, Brightly Fluorescent, L5 Pyramidal Neuron Dendrites with Clearly Resolvable Dendritic Spines.**

**Supplementary Movie 2. Example Rotarod Performance in an Imaged *MECP2*- duplication Mouse (right) and a Wild-type Littermate (left).** Note the markedly better rotarod performance of the mutant. Movie is accelerated 2 times.

